# A modular mathematical model of exercise-induced changes in metabolism, signaling, and gene expression in human skeletal muscle

**DOI:** 10.1101/2021.05.31.446385

**Authors:** I.R. Akberdin, I.N. Kiselev, S.S. Pintus, R.N. Sharipov, A.Yu. Vertyshev, O.L. Vinogradova, D.V. Popov, F.A. Kolpakov

## Abstract

Skeletal muscle is the principal contributor to exercise-induced changes in human metabolism. Strikingly, although it has been demonstrated that a lot of metabolites accumulating in blood and human skeletal muscle during an exercise activate different signaling pathways and induce expression of many genes in working muscle fibres, the system understanding of signaling-metabolic pathways interrelations with downstream genetic regulation in the skeletal muscle is still elusive. Herein, a physiologically based computational model of skeletal muscle comprising energy metabolism, Ca^2+^ and AMPK signalling pathways, and expression regulation of genes with early and delayed responses has been developed based on a modular modeling approach. The integrated modular model validated on diverse including original experimental data and different exercise modes provides a comprehensive *in silico* platform in order to decipher and track cause-effect relationships between metabolic, signaling and gene expression levels in the skeletal muscle.

## 1 Introduction

Skeletal muscle tissue comprises about 40 % of total body mass in adult lean humans and plays a crucial role in the control of whole-body metabolism and exercise tolerance. Regular low-intensity exercise (aerobic or endurance training) strongly increases vascular and mitochondrial density, oxidative capacity, improving fat and carbohydrate metabolism. These adaptations lead to an enhancement of muscle endurance performance and reduce the risk associated with the morbidity and premature mortality of chronic cardiovascular and metabolic diseases (Pedersen & Febbraio, 2012; Hawley et al., 2014).

Acute aerobic exercise induces significant metabolic changes in the working skeletal muscle which in turn activate numerous signaling molecules. Changes in the content of Ca^2+^ ions in skeletal muscle plays a fundamental role in the regulation of the activity of contractile proteins and enzymes involved in energy metabolism. In addition, a contractions-induced increase in the content of Ca^2+^ ions in the myoplasm significantly affects the activation of some signaling proteins: Ca^2+^/calmodulin-dependent kinases (CaMKs), calcineurin, Ca^2+^-dependent protein kinase C, etc. (Koulmann & Bigard, 2006). Increasing the intensity of contractile activity more than 50% of maximal pulmonary O_2_ consumption rate (V’O2max) induce a linear increase in the activity of AMPK (AMP-dependent protein kinase), the key energy sensor of the cell activated by an increase in the AMP/ATP ratio, Ca^2+^ -dependent kinase CaMKKII and a decrease in the level of muscle glycogen (Richter & Ruderman, 2009). In muscle cells, activated AMPK changes the level of phosphorylation of several dozen different signaling proteins (Hoffman et al., 2015; Needham et al., 2019; Nelson et al., 2019). Thus, Ca^2+^ and AMPK play a key role in the regulation of various intracellular signaling cascades, as well as the gene expression induced by an exercise.

Dramatic changes in the expression of hundreds genes were observed during the first hours of recovery after acute intensive aerobic exercise in untrained skeletal muscle (Vissing & Schjerling, 2014; Dickinson et al., 2018; Popov et al., 2018) as well as in muscle adapted to regular exercise training (Neubauer et al., 2013; Vissing & Schjerling, 2014; Popov et al., 2018). These changes are associated with muscle contraction *per se* and with system factors and circadian rhythms. On the basis of the analysis of differentially expressed genes between exercised and contralateral non-exercised vastus lateralis muscle, the contractile activity-specific transcriptome responses at 1 and 4 hours after the one-legged exercise were identified in our previous study (Popov et al., 2019). It was shown that the most enriched biological process for the transcriptome response is transcription regulation, i.e. increase in expression of genes encoding transcription factors and co-activators. The study demonstrated that genes encoding such transcription factors as *NR4A, AP-1* and *EGR1* were actively expressed at 1 hour after the termination of the exercise, while other transcription regulators like *PPARGC1A, ESRRG* and *VGLL2* were highly expressed at 4 hour. Both sets of transcription factors modulate muscle metabolism. We suggest that gene expression on early and late stages of the recovery after the termination of the exercise can be regulated by different ways. Obviously, these molecular mechanisms are pretty complex, but we suppose that the expression of genes with early and delayed response to a stress stimulus may be ensured by means of some general or basic mechanisms of the gene expression regulation.

It is worth to note, although advancement in the development of high-throughput experimental techniques and generation of diverse omics data for human skeletal muscle during endurance exercise enabled to unveil key participants of the cellular response and adaptation to stress/various stimuli (Neubauer et al., 2013; Vissing & Schjerling, 2014; Dickinson et al., 2018; Popov et al., 2018; Popov et al., 2019), the system understanding of signaling-metabolic pathways relationships with downstream genetic regulation in exercising skeletal muscle is still elusive. Detailed mechanistic and multiscale mathematical models have been constructed to provide a powerful *in silico* tool enabling quantitatively investigating activation of metabolic pathways during an exercise in skeletal muscle (Li et al., 2009; Akberdin et al., 2013; Kiselev et al., 2019). Here, we propose a modular model of exercise-induced changes in metabolism, signaling and gene expression in human skeletal muscle. The model includes different compartments (blood, muscle fibers with cytosol and mitochondria) and allows ones to quantitatively interrogate dynamic changes of metabolic and Ca^2+^- and AMPK-dependent signaling pathways in response to an aerobic exercise of various intensity in slow- and fast-twitch muscle fibres (type I and II, respectively), as well as downstream regulation of genes with early and delayed response in a whole/mixed fiber type skeletal muscle.

The model modules are hierarchically organized and present accordingly metabolic, signalling and gene expression levels. To build the model we have used the BioUML platform (Kolpakov et al., 2019) that is designed for modular modelling of complex biological systems. The effectiveness of both this approach and BioUML platform was previously confirmed by the development of complex modular models of apoptosis (Kutumova et al., 2012) and cardiovascular system (Kiselev et., 2012).

## 2 Methods

### Model construction

#### BioUML platform for model reconstruction

BioUML (Biological Universal Modeling Language, https://ict.biouml.org/) (Kolpakov et al., 2019) is an integrated Java-based platform for modeling biological systems. The tool is applicable to solve a wide range of tasks including access to diverse biological databases, mathematical description and visual representation of biological systems, numerical calculations and parametric and other types of model analysis. The main features of the BioUML are:

- ability to use both standalone version of the tool and web-services remotely;
- support of generally accepted standards for model description of biological systems (SBML) and their graphical notation (SBGN) (Le Novere et al., 2009; Hucka et al., 2018);
- visual modeling of biological systems and processes: a user has an opportunity to graphically construct and edit the developed model;
- support of different mathematical representations (ordinary differential equations, algebraic equations, discrete events and stochastic modeling);
- modular platform architecture that facilitates extension and/or addition of new types of models, methods of numerical calculations, etc.

The BioUML platform allows users to clone, modify and store multiple versions of each module in the repository. It ensures flexibility of a modular model comprising individual modules by means of a combination of different versions of modules for diverse purposes.

#### Visual modeling

Representation of investigating systems as graphical diagrams by means of a software supporting visual modeling can significantly facilitate procedure of the model reconstruction. We consider visual modeling as a formal graphical representation of the system and/or modeling processes as a diagram and consequent dynamic simulation based on the representation. Graphical notation is a crucial component of visual modeling which allows one to formally and completely build a model. A visual model can be presented by some types of diagrams enabling description of diverse aspects of the structure and function of a complex system with different levels of details. This formal graphical representation is a basis for automatic code generation by specialized tools to simulate the model. We have made an extension of well-known SBGN notation (Le Novere et al., 2009) in order to simulate physiological processes which require not only description between metabolites, but also ability to use algebraic, differential equations and instant transition of the system from one state to another. In addition, connections between equations indicate signal transduction in the model while interface ports of modules (or submodels) have also a direction (input, output or contact). A meta-model is a basis of the visual modeling in BioUML which ensures a formalism for complex description, graphical representation and numerical simulation of biological systems on different levels of their hierarchical organization. A meta-model consists of three interrelated levels of complex systems description:

- graphical representation - system’s structure is described as a compartmentalized graph;
- database - each element of the graph can include reference on a certain object in the database;
- runned model - an element of the model (variables, mathematical equations, discrete events, states and transitions) can be associated with an element of the graph (vertices, arcs and compartments). As an example, vertices of the graph can be represented by variables or states of the system, while arcs of the graph correspond to equations describing changes of these variables or transitions between two states.

The description of the biological system as a meta-model is used to generate a Java code reflecting the model as a system of algebraic and/or differential equations, considering delay components, piecewise functions, discrete events and transitions. To generate a code the specific simulation engine is employed which defines the model type and corresponding simulation method.

#### A multi-compartmental complex model

A BioUML diagram describing a modular multi-compartmental model contains interconnected elements or modules (submodels) each of which is referred to another diagram (also may be modular). A directed connection between input and output nodes determines the signal transduction from a module to another one, while undirected relation between contacts reflects signals exchange between modules (Figure 1). According to this methodology, an integrated mathematical model describing energy metabolism of the human skeletal muscle (Kiselev et al., 2019; Li et al., 2012) considering Ca^2+^- and AMPK-dependent signaling pathways and downstream regulatory processes of the expression of early response and genes with delayed response has been built. A complex mathematical model (Li et al., 2012) developed by Li and coathours in MATLAB (https://www.physiome.org/jsim/models/webmodel/NSR/Li2012/) has been rebuilt in BioUML as an initial model of the energy metabolism of the human skeletal muscle taking into account quantitative differences between fiber I and II types (Figure 2). An activation mechanism which enhances energy metabolism via transport and reaction fluxes due to physical exercise was harnessed as the stress function depending on general work rate parameter. The work rate parameter defines power of the physical exercise and variates depending on the mode of the exercise. In our model muscle volume was 5 L that corresponds to that involved in exercise at cycling ergometer.

**Fig. 1.**
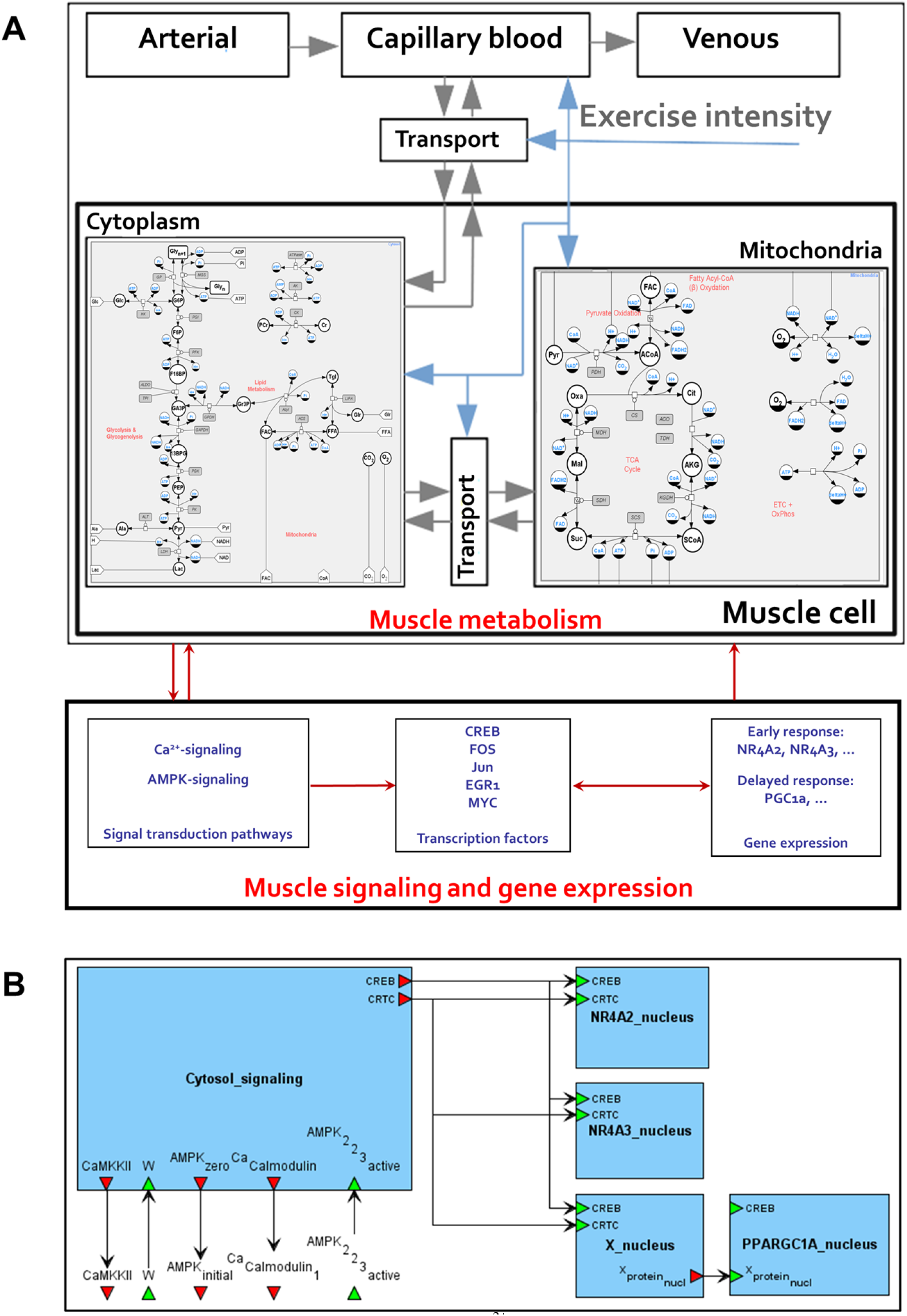
An integrated modular model linking metabolism, Ca^2+^-dependent signaling transduction and regulation of gene expression in human skeletal muscle. Grey arrows in the metabolic module represent transport reactions of metabolites, blue arrows mean external control function like consideration of physical exercise. Module «Muscle tissue» consists of submodules «Cytosol» and «Mitochondria», which in turn contain equations describing enzymatic reactions inside the certain compartment (A). A certain mathematical module has been developed for genes with early and delayed response (B).

**Fig. 2.**
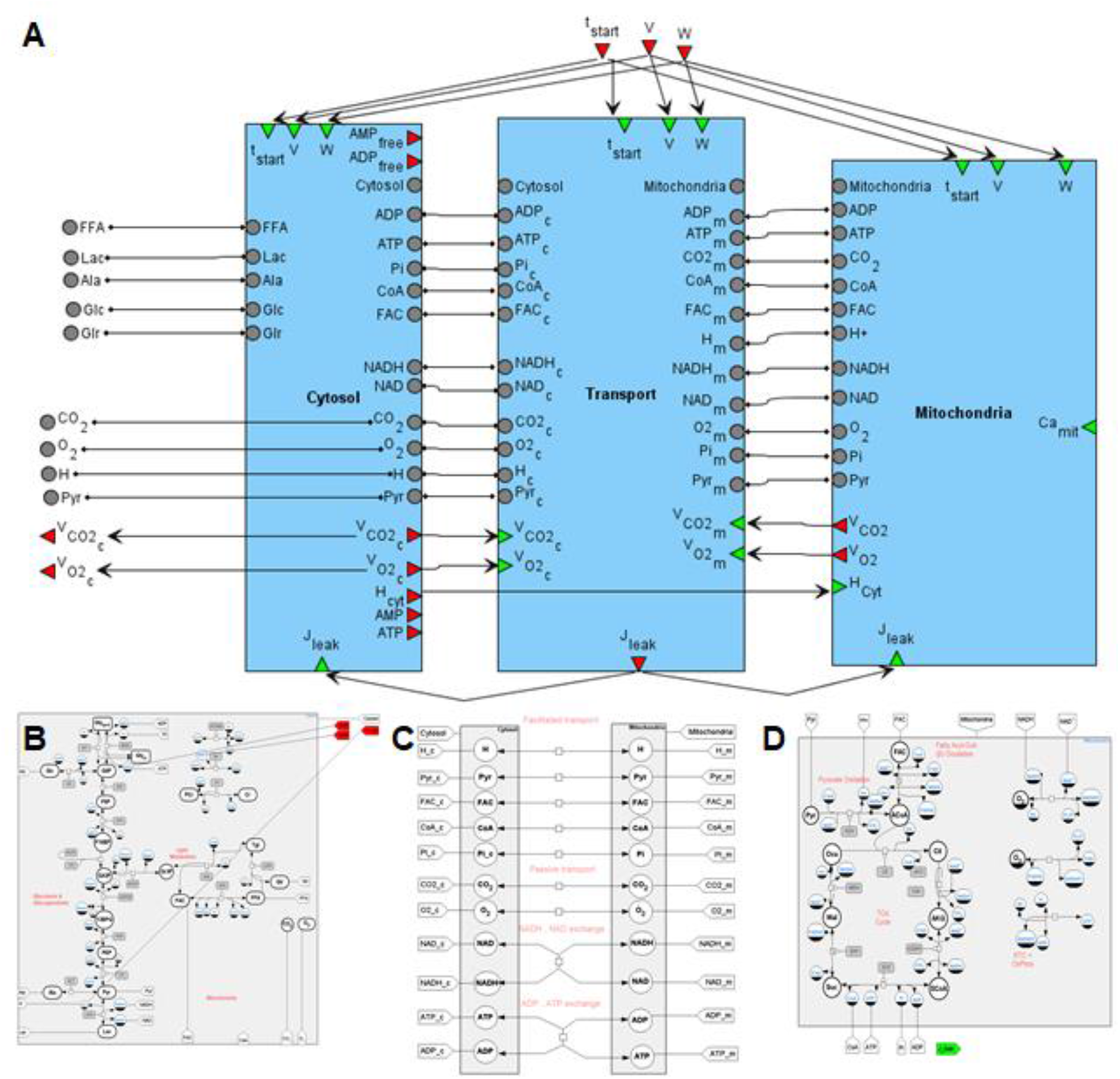
A general SBGN diagram of the modular model describing metabolism in human muscle fibers (A) taking into account metabolic processes in the cytoplasm (B), in the mitochondrion (C) and transport reactions between two compartments (D).

#### Upgrade of the model (metabolic level)

It is worth noting that values of activation coefficients associated with ATPase (Stienen et al., 1996; He et al., 2000; Szentesi et al., 2001; Barclay, 2017) and pyruvate dehydrogenase reaction fluxes for type I and type II fibers (Parolin et al., 1999; Kiilerich et al., 2008; Albers et al., 2015) as well as time constant of ATPase flux rate coefficient in response to exercise were modified (See data availability) according to recently published data and estimations (Broxterman et al., 2017; Bartlett et al., 2020).

Despite overall net glycogen breakdowns during muscle contraction, exercise increases the activity of glycogen synthase (GS) (Wojtaszewski et al., 2001; Nielsen & Richter, 2003; Jensen et al., 2009; Jensen & Richter, 2012) and ATP consumption related with the reaction. Therefore, GS reaction fluxes were modified according to (Wojtaszewski et al., 2001; Jensen et al., 2009; Jensen et al., 2012b). The rates of muscle glycogen synthesis during exercise assumed to be equal in type I and type II fibres and were estimated from average post-exercise glycogen synthesis data (Casey et al., 1995).

To consider the allosteric regulation of AMPK activity (in corresponding modules, Fig. 1, data availability) concentrations of free ADP and AMP in the cytosol were calculated using intracellular Cr, PCr, ATP and H^+^ concentrations as well as the equilibrium constants for creatine phosphokinase and adenylate kinases in each fiber type as described previously (Lawson & Veech, 1979; Dudley & Terjung, 1985; Mannion et al., 1993) (Figure 3).

**Fig. 3.**
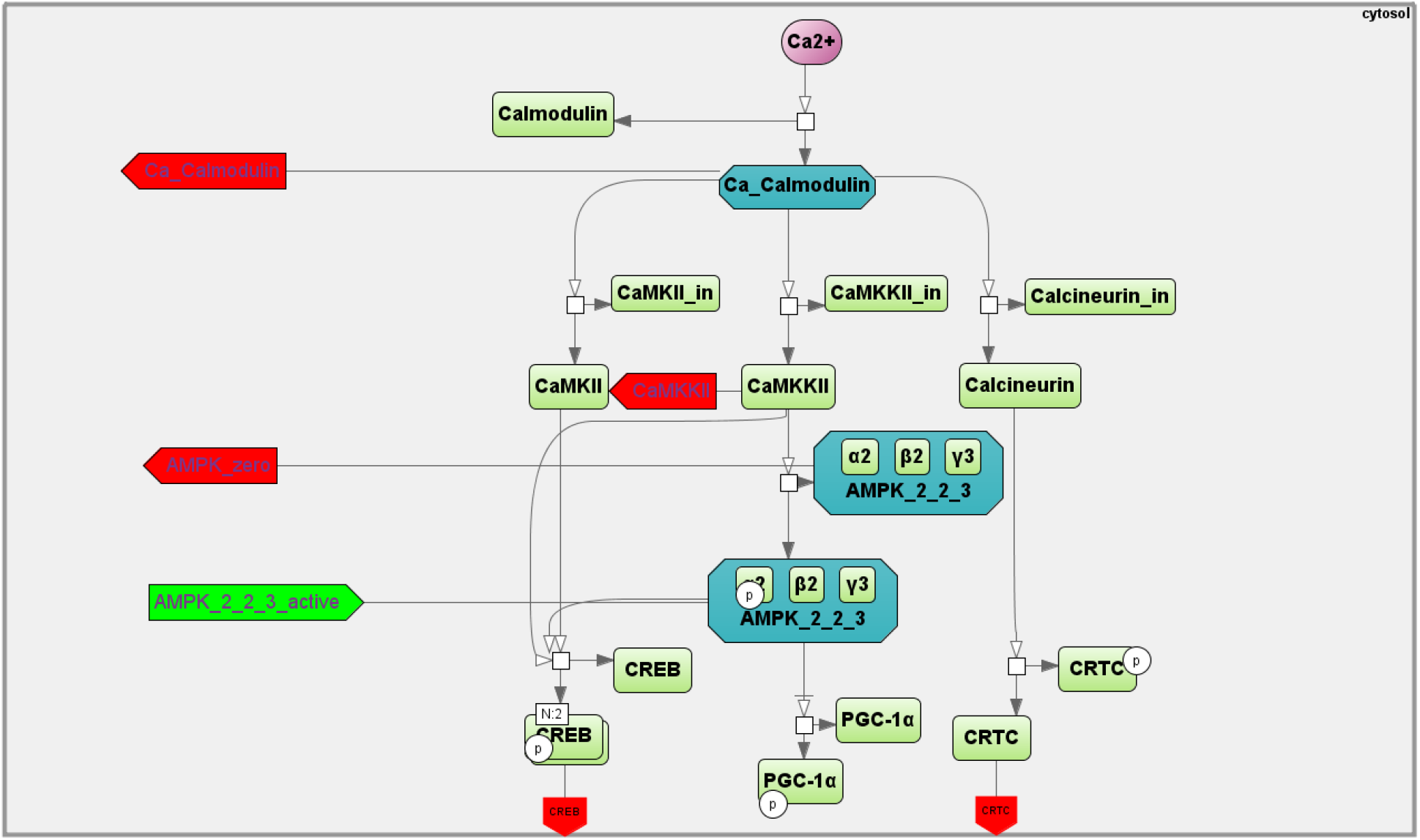
A SBGN diagram of the Ca^2+^- and AMPK-dependent signaling pathway activating by contractile activity (aerobic exercise).

The diagram of the modular model describing metabolism of human skeletal muscle is presented on Figure 2. The cytosol includes metabolic reactions of the glycolysis, glycogenolysis and lipids metabolism, while tricarboxylic acid (TCA) cycle, ß-oxidation and oxidative phosphorylation reactions are presented in the mitochondria. The intermediate compartment between those is a transport module which contains passive and facilitated transport reactions for model intracellular species. Kinetic laws presenting metabolic and transport flux expressions exactly match the initial model developed by Li and coauthors (Li et al., 2012).

According to the model a dynamic mass balance of metabolites (*i*) is based on the structural and functional organization and is expressed by equations:

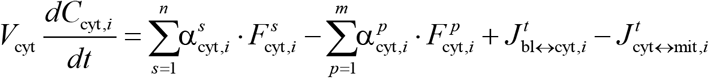

in the cytosol and

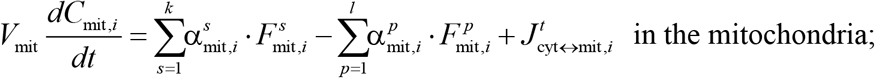

where *V*_*cyt*_, *V*_*mit*_ is a volume of the corresponding module or compartment; *C*_*cyt,i*_, *C*_*mit,i*_ is a concentration of the metabolite *i* in a certain compartment; 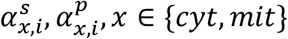 are corresponding stoichiometric coefficients in the metabolite consumption or production reactions in a certain compartment; 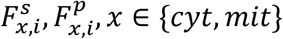 are corresponding velocity laws of the metabolite consumption or production reactions in a certain compartment; 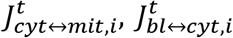 are transport fluxes between cytosol and mitochondria compartments, cytosol and blood compartments, correspondingly.

In order to describe a dynamic mass balance of metabolites (*i*) in the blood compartment the next equation is used:

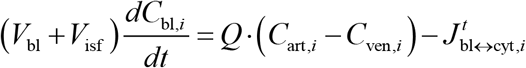

where *V*_*bl*_, *V*_*isf*_ is a volume of the capillary blood and interstitial compartment, correspondingly; *C*_*art,i*_, *C*_*ven,i*_, *C*_*bl,i*_ is a concentration of the metabolite *i* in arterial, venous and capillary blood compartments, correspondingly; *Q* is a blood flow.

It is worth to note that such modules as capillary blood and interstitial fluid are assumed to be in equilibrium with each other that allows us to consider species concentrations in these compartments as equal and combine both of them into the blood compartment.

#### Signaling level

The mean concentration of Ca^2+^ ions in the myoplasm increases in proportion to intensity of exercise. Ca^2+^ binds to calmodulin, thereby activating CaMKs and phosphatase calcineurin (Gehlert et al., 2015). CaMKII is the most abundant isoform in the human skeletal muscle, whereas CaMKI and CaMKIV are not expressed at detectable levels (Rose et al., 2006). An increase in CaMKII activity results in CREB1 Ser133 phosphorylation (and likely some other CREB-like factors) leading to activation of the transcription factor (Johannessen & Moens, 2007; Olesen et al., 2010). Calcineurin can dephosphorylate (and activate) CRTCs at Ser171 (CREB-regulated transcription coactivators) playing a key role in regulating transcriptional activity of CREB1 (Altarejos & Montminy, 2011). Another target of calmodulin is Ca^2+^/calmodulin-dependent protein kinase kinase 2 (CAMKK2) that phosphorylates AMPK Thr172 thereby activating the kinase (Abbott et al., 2009). In turn, activated AMPK can phosphorylate CREB1 Ser133 (Thomson et al., 2008). Collectively, these findings drove us to include in our model the Ca^2+^-dependent regulation of calmodulin, CREB1 (via CaMKII), CRTC (via calcineurin), and AMPK (via CaMKK2) (Figure 3). The amount of these proteins in human skeletal muscle was estimated using published proteomics and transcriptomics data (Murgia et al., 2017; Popov et al., 2019) (<u>see Supplementary data in</u> (Akberdin et al., 2020)).

There are three different heterotrimeric complexes of AMPK in the human skeletal muscles: α2β2γ1, α2β2γ3, and α1β2γ1 (Wojtaszewski et al., 2005). Distinct kinetic properties (an intrinsic enzyme activity, binding affinities of AMP, ADP and ATP to the specific isoform, sensitivity to de- and phosphorylation of AMPK heterotrimers) (Rajamohan et al., 2016; Ross et al., 2016) and their subcellular localization (Pinter et al., 2013) cause a differential regulation of the AMPK heterotrimers *in vivo*. The α2β2γ3 complex is phosphorylated and activated during moderate- to high-intensity aerobic exercise, while the activity associated with the other two AMPK heterotrimers is almost unchanged (Birk & Wojtaszewski, 2006). However, the baseline activity of the α2β2γ3 complex is significantly lower than others. The general AMPK activity in baseline and during/after exercise is a sum of isoforms activities, hence, we considered separately all isoforms (in the corresponding module) to quantitatively fit an experimental data obtained at baseline and after an exercise (Birk & Wojtaszewski, 2006; Willows et al., 2017). AMPK is regulated by various ways: an up-stream kinase LKB1 can phosphorylate AMPK at Thr172 (Lizcano et al., 2004; Jansen et al., 2009). On the other hand, an exercise-induced decrease in intramuscular ATP increases its activity, while an increase in AMP activates it (Hardie, 2016; Li et al., 2017). Hence, in our model the AMPK is regulated via AMP, ATP, and LKB1, as well as CaMKK2 (as mentioned above) (Figure 3).

#### Gene expression level

An aerobic exercise induces expression of several hundreds of genes regulating many cell functions: energy metabolism, transport of various substances, angiogenesis, mitochondrial biogenesis, etc. Regulation of the transcriptomic response to acute exercise includes dozens of transcription regulators (Popov et al., 2019) and seems to be extremely complex. Therefore to consider the response on gene expression level we exemplify the regulation of some genes encoding the nuclear receptors NR4As and a transcription co-activator PGC-1α (encoded by *PPARGC1A* gene) -key exercise-induced regulators of the angiogenesis, mitochondrial biogenesis, fat and carbohydrate metabolism in skeletal muscle (Lira et al., 2010; Pearen & Muscat, 2018).

Expression of *NR4A2, NR4A3* mRNA rapidly increases during the first hour after an aerobic exercise (early response genes) (Popov et al., 2019) due activation of Ca^2+^\calcineurin-dependent signaling (Pearen & Muscat, 2018). We included in our model the Ca^2+^-dependent regulation (Ca^2+^\calcineurin-CaMKII-CREB1) of *NR4As* genes using data of contractile activity-specific mRNA response of these genes (Popov et al., 2019). Expression of *PPARGC1A* mRNA rises 3 to 4 h after an exercise (gene with delayed response) (Popov et al., 2019). The transcription regulation of *PPARGC1A* via the canonical (proximal) and inducible (distal) promoters is very complicated, and includes Ca^2+^- and AMPK-dependent signaling, as well as CREB1 and its co-activator CRTC (Popov et al., 2015; Popov et al., 2018). The phosphorylation level of many signaling kinases drops to basal levels within the first hour after an aerobic exercise. Moreover, in a genome-wide study on various human tissues, it was shown that the phosphorylation level of CREB Ser133 does not always correlate with its transcriptional activity (Zhang et al., 2005). Therefore, we suggested the expression of genes with delayed response (including *PPARGC1A*) is regulated by increasing the expression of one of the early response genes encoding transcription factors leading to a rapid increase of corresponding protein (see Figure 1 and Figure 5 in (Akberdin et al., 2020). Analysis of contractile activity-specific transcriptomic data (Popov et al., 2019) showed that a rapid increase in the expression of genes encoding various TFs is observed already in the first hour after an exercise. It turned out that the binding motifs of some TFs (CREB-like proteins, as well as proteins of the AP-1 family: FOS and JUN) are located and intersected with each other both in the alternative and in the canonical promoters of the *PPARGC1A* gene (Akberdin et al., 2020), i.e. these TFs can act as potential regulators of this gene. This is consistent with the fact that these TFs can bind to DNA and regulate the expression of target genes as homo- and heterodimers (Hai & Curran, 1991; Newman & Keating, 2003). Based on these considerations, we included in the model the regulation of gene expression of early (*NR4A2, NR4A3*) and delayed (*PPARGC1A*) genes: early response genes are regulated via the activation of existent TFs (e.g. CREB1) and their co-activators (e.g. CRTC), while delayed response genes - via an increase in the expression of early response genes encoding transcription factors (transcription factor X in our model, Fig. 1).

### Model simulations

Numerical solutions of the model represented by a system of ordinary differential equations have been obtained on the basis of the VODE method (Brown et al., 1989) using a JVode simulation engine implemented in BioUML tool (Kolpakov et al., 2019). Each submodule of the modular model can be represented as an independent SBML file (Hucka et al., 2018), while the integrated modular model can be exported as a COMBINE archive (Bergmann et al., 2014) in order to use the model and reproduce simulations results in alternative software supporting current standards of the systems biology.

## 3 Results

### Model verification

#### Simulation of metabolic changes induced by incremental and interval exercises

To validate the metabolic part of the model we investigated dynamic behaviour of the system in response to diverse acute aerobic exercises and compared them with published experimental data. Initially, we quantitatively estimated biochemical responses of the key metabolic variables (ATP, ADP, PCr, lactate concentrations and pH in muscle fibers type I and II) in the incremental ramp exercise to exhaustion which is a commonly used approach to evaluate aerobic performance. Increasing the power during the ramp exercise effects on various physiological variables like the number/volume of recruited muscle fibre type I and II, blood flow as well as of transport and metabolic fluxes in both fibre types (Fig. 4 and see data availability). In our simulation, muscle fibres type I are started to be recruited after the beginning of exercise, while fibre type II - only at power over 24% of VO2_max_ (6 min after the ramp exercise onset, Fig. 4A). Recruiting all muscle fibres during the test lead to exhaustion and termination of the exercise (Bottinelli et al., 1996; Stienen et al. 1996; Li & Larsson 2010); the peak power at exhaustion in our simulation was 250 W that corresponds to the value for untrained male performing the ramp exercise till exhaustion at cycling ergometer. The model simulations reasonably well correspond to experimental measurements (Roussel et al., 2003; Greiner et al., 2007; Cannon et al., 2013) obtained in studies with the incremental exercise (Supplementary Figure1). It is worth to note that the current version of the model does not take into account an effect of the muscle fatigue during the incremental ramp exercise observed in exercised muscle *in vivo* (see below). The fact may partially explain the lack of exponential changes in muscle lactate concentration and pH during the last part of the incremental exercise.

**Fig. 4.**
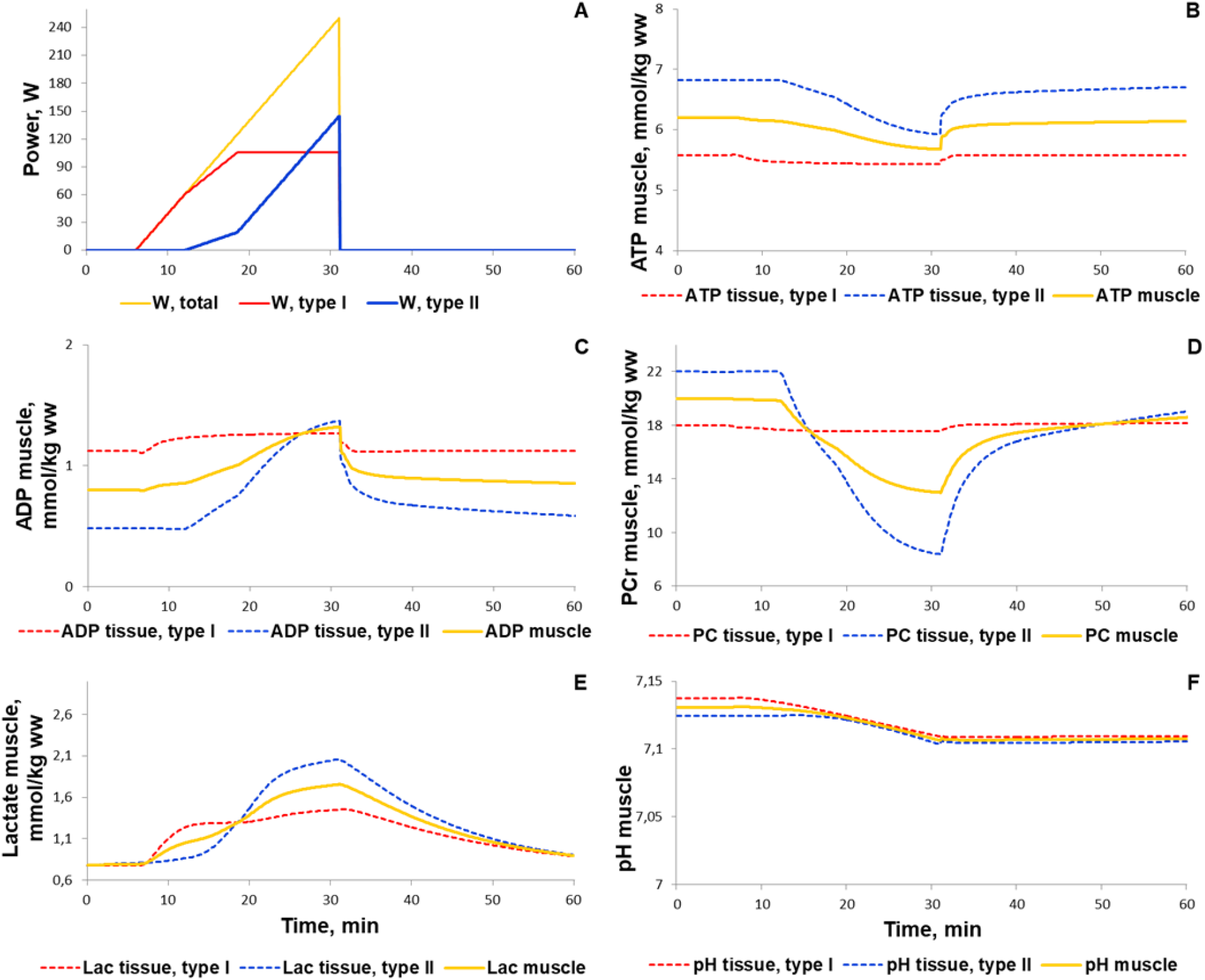
Simulation results for the incremental ramp exercise till exhaustion. (A) Exercise power and fibers recruitment pattern: total power (orange), power generated by type I (red) and II (blue) fibers; (B, C, and D) ATP,ADP, and PCr concentrations in type I (red, dotted) and II (blue, dotted) fibers and in the muscle tissue (orange, solid); (E and F) Lactate concentrations and pH changes in type I (red, dotted) and II (blue, dotted) fibers and in the muscle tissue (orange, solid).

For additional verification of the metabolic part of the model we simulated responses to various interval exercises (Fig. 5, 6). Figure 5 shows that the model qualitatively reproduces the dynamics of PCr concentration during the interval exercises with different patterns (duration of an exercise bout 16 s to 64 s and recovery period 32 s to 128 s) and with peak power comparable with maximal aerobic power obtained in the incremental ramp test (250 W) (Davies et al., 2017).

**Fig. 5.**
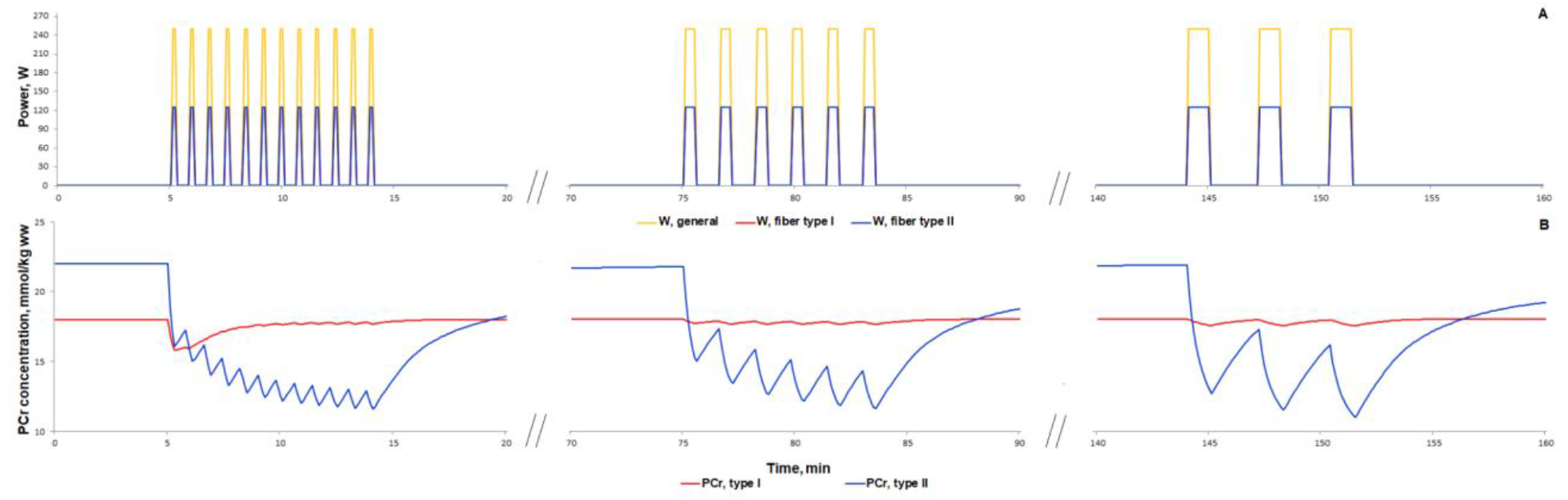
Simulation results for high-intensity intermittent exercise bouts with different ratios of work:recovery (initial - 16:32 s; intermediate - 32:64 s; final - 64:128 s) (Davies et al., 2017). (A) Exercise power and fibers recruitment pattern: total power (W_peak_=250) (orange), power generated by type I (red, W_peak_=125) and II (blue, W_peak_=125) fibers; (B) PCr concentration in type I (red) and type II fibers (blue).

**Fig. 6.**
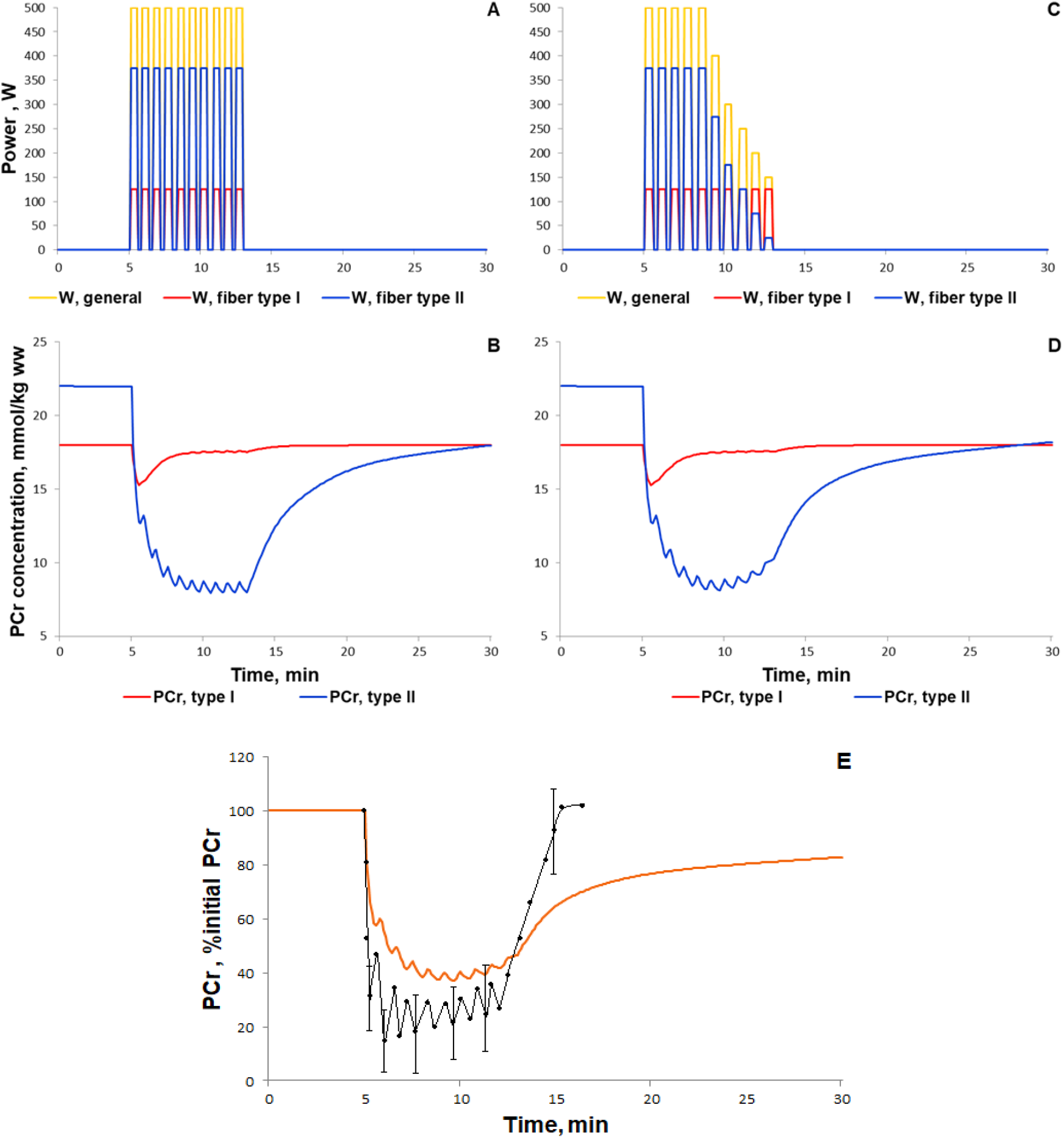
Simulation results for high-intensity intermittent exercise (each bout of 30-s exercise separated by 20-s recovery; Kappenstein et al., 2013). (A, C) Exercise power and fibers recruitment pattern: total power (A: W_peak_=500; C: W_peak_=500), W (orange), power generated by type I (red, A-C: W_peak_=125) and II (blue, A: W_peak_=375; C: W_peak_=375 and successive power decline) fibers; (B, D) PCr concentration in type I (red) and type II fibers (blue); (E) Changes in PCr concentration (% initial): orange line is the simulation result for PCr in the muscle tissue, while black dots with corresponding line is the experimental data from Kappenstein et al., 2013 (mean ± SD for some dots).

High intensity interval exercise has been shown to rapidly decrease in the PCr level followed by slow recovery of the PCr concentration during the last part of the exercise (Kappenstein et al., 2013). There is no data concerning the exercise power in the study hence we used the constant value (500 W) for each bout (Fig. 6A). The power is markedly greater than the peak power in the incremental cycling test because the duration of each exercise bout is short; the energy supply of such short exercise bouts is related mainly with PCr reactions as well as glycolysis. Our model precisely simulated the rapid decline in PCr, but showed no slow recovery of the PCr during the last part of the high intensity interval exercise (Fig. 6B). We suggested that this discrepancy may be related with the lack of the fatigue-induced decline in exercise power. We tried to roughly simulate the fatigue-induced decline in exercise power by the decline in power generated by muscle fibers type II (Fig. 6C). As a result, the model much better reproduced the experimental dynamics of PCr than simulations with constant maximal power in each bout (Fig. 6D, E). However, PCr dynamics during the recovery process indicates that the model still requires further modifications and numerical study. We suppose that the potential point for the update is related to the pH changes during the recovery.

#### Simulation of signaling and gene expression changes induced by low- and moderate intensity continuous exercises

At the next step of the model validation we predicted the responses of biochemical variables, signaling molecules (AMPK and Ca^2+^-dependent proteins), transcription factor (CREB1), as well as expression of genes with early and delayed response (*NR4A3, NR4A2, PPARGC1A*) to low (50% VO2_max_) and moderate intensity (70% VO2_max_) continuous aerobic exercises. Moderate intensity exercise recruits more muscle fibers type II than low intensity exercise thereby additionally modulating the exercise-induced metabolic fluxes and molecular response. Comparison of our simulations with experimental data (Krustrup et al., 2004; Barker et al., 2008; Cannon et al., 2014; Fiedler et al., 2016; Bartlett et al., 2020) showed that the model reproduces well the metabolic changes in various fiber types and in the whole muscle induced by exercises with various intensity (Fig. 7; Supplementary Figure 1).

**Fig. 7.**
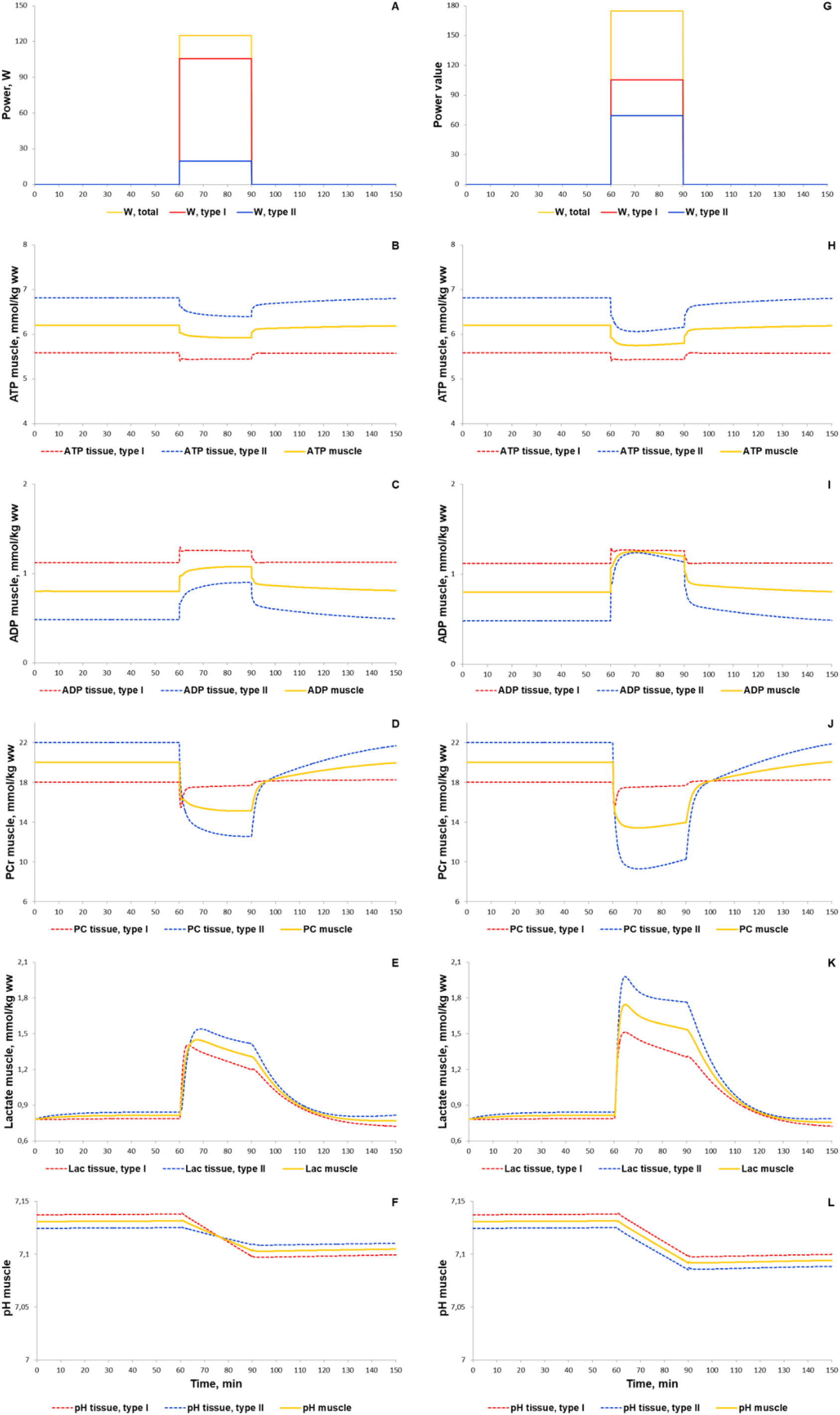
Simulation results for low (50% VO2_max_, A-F) and moderate intensity (70% VO2_max_, G-L) continuous exercises (30 minutes). (A, G) Exercise power and fibers recruitment pattern: total power (orange), power generated by type I (red) and II (blue) fibers; (B, H) ATP, ADP, PCr, lactate concentrations and pH changes in type I (red, dotted) and II (blue, dotted) fibers and in the muscle tissue (orange, solid) during low (B-F) and moderate (H-L) intensity exercise.

According to the literature data on the human *vastus lateralis* muscle (Birk & Wojtaszewski, 2006; Rose et al., 2006; Lee-Young et al., 2009), our simulation shows an intensity-dependent increase in phosphorylation of CAMKII and AMPK α2 and γ3 (Fig. 8 A - C, F - H). Importantly, the phosphorylation (as a marker of activity) of AMPK α2 and γ3 consists of 10% and 30% of the AMPK isoforms containing α2 and γ3, respectively (Fig. 8 B-C, G-H), that is in line with the experimental data (Birk & Wojtaszewski, 2006). Unlike experimental data on signaling level, we found transcriptomics data concerning intensity-dependent gene expression for 1 hour exercise only (Popov et al., 2018; 2019). In our model, exercise-induced activation of CAMKII and AMPK induces CREB- and CRTC-related expression of early response genes that is in line with the experimental data (Popov et al., 2019) on exercise-induced expression of early response genes (by example of *NR4A2, NR4A3*) in the human *vastus lateralis* muscle (Fig. 8K).

**Fig. 8.**
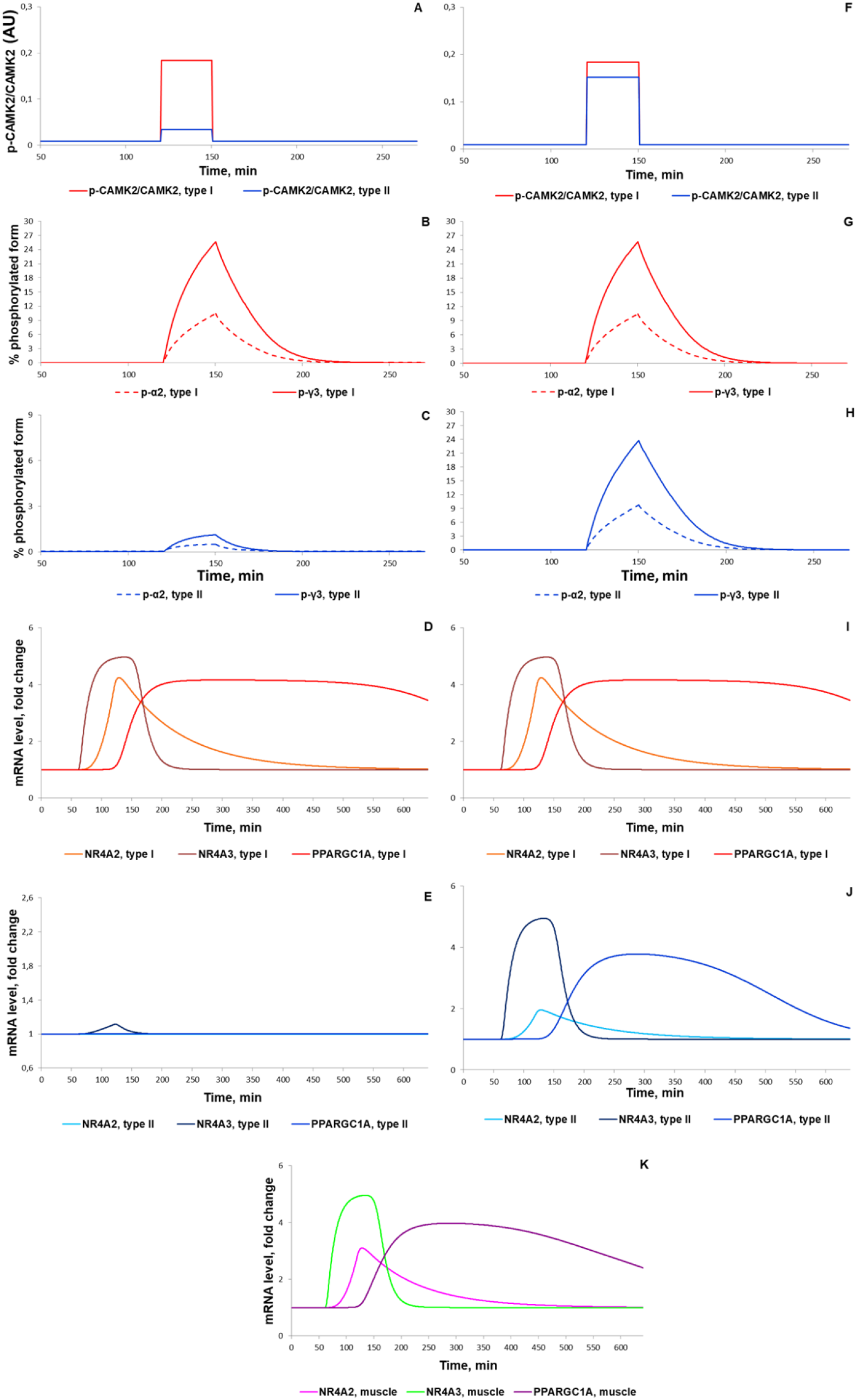
Simulation results for low and moderate intensity (50% (A-E) and 70% (F-K) VO2_max_, respectively) continuous exercises (30 minutes for signaling and 60 min for gene expression) with intermediate X factor regulating expression of *PPARGC1A* gene. (A,F) Ratio of CAMKII phosphorylated protein in type I (red) and type II (blue) fibers (corresponds to Rose et al., 2006); (B, G, C, H) Percentage of all α2 phosphorylated protein (dashed) and of the phosphorylated γ3 heterotrimers (solid) in type I fibers and type II fibers, respectively (corresponds to Birk & Wojtaszewski, 2006; Lee-Young et al., 2009); (D, I) Expression (in fold changes) of *NR4A3* (thunderbird solid), *NR4A2* (orange solid), *PPARGC1A* (red solid) in type I fibers and (E, J) in type II fibers, where *NR4A3* - sapphire solid, *NR4A2* - azure solid, *PPARGC1A* - blue solid; (K) Expression (in fold changes) of *NR4A3* (green solid), *NR4A2* (magenta solid), *PPARGC1A* (purple solid) in the muscle tissue for moderate intensity exercise.

Numerical analysis of the model demonstrated the necessity of consideration of additional transcription factors showing activity at 1 to 2 h after exercise for simulation of genes with delayed response to exercise (by example, *PPARGC1A*; Akberdin et al., 2020). Introducing in the model transcription factor X that expression is up-regulated immediately after exercise in CREB- and CRTC-dependent manner allowed to reproduce expression of *PPARGC1A* gene (Fig. 8K). Our bioinformatics analysis (Akberdin et al., 2020) of the transcriptomics data (Popov et al., 2019), in turn, allowed to suggest that proteins from the AP-1 family (e.g. FOS and JUN) forming heterodimer complexes with CREB-like transcription factors are served as these intermediate regulators (factor X; Fig. 1). Moreover, an analysis of the transcriptomic (Catoire et al., 2012; Popov et al., 2019) and ChIP-seq data from the GTRD database (Yevshin et al., 2019) revealed that expression of *PPARGC1A* via the alternative promoter may be regulated by EGR1 and MYC. Both *EGR1* and *MYC* markedly induce expression in 30 to 60 minutes after an aerobic exercise and have binding motifs in the alternative promoter. Our prediction is supported by experimental data showing the *EGR1* expression leads to an increase of *PPARGC1A* expression in human aortic smooth muscle cells. (Fu et al., 2002; 2003), while the *EGR1* expression promptly and dramatically increases after stretching of skeletal muscle cells leading to an increase in the concentration of EGR1 protein in 3-4 hours (Pardo et al., 2011). On the one hand, MYC positively regulates the expression of all active genes through transcriptional amplification (Lin et al., 2011; Nie et al., 2012; Rahl & Young, 2014) and chromatin modifications (Guccione et al., 2006; Knoepfler et al., 2006). However, an enhancement of its expression negatively impacts *PPARGC1A* expression (Tan et al., 2016; Mastropasqua et al., 2018), in particular, in cardiomyocytes (Ahuja et al., 2010) and other types of cells where MYC acts as repressor (Sancho et al., 2015).

## 4 Discussion

### The Integrated modular model comprises three hierarchical levels (metabolic, signaling and gene expression)

We have previously developed a multi-compartmental mathematical model describing the dynamics of intracellular species concentrations and fluxes in human muscle at rest and intracellular metabolic rearrangements in exercising skeletal muscles during an aerobic exercise on a cycle ergometer (Kiselev et al., 2019). As an initial model for this study, we have used a complex model of energy metabolism in the human skeletal muscle developed by Li and coauthors and considered two types of the muscle fibers (Li et al., 2012). We have proposed a modular representation of the complex model using BioUML platform (Kolpakov et al., 2019). The modular representation provides the possibility of rapid expansion and modification of the model compartments to account for the complex organization of muscle cells and the limitations of the rate of diffusion of metabolites between intracellular compartments. This feature allowed us to integrate modules of signalling pathways modulating downstream regulatory processes of early response and genes with delayed response during the exercise and recovery. The parametric fitting of the modular model to published experimental data (Krustrup et al., 2004; Greiner et al., 2007; Barker et al., 2008; Cannon et al., 2014; Bartlett et al., 2020; see Supplementary Figure 1) showed the validity of the modular modeling approach implemented in BioUML. Furthermore, the integrated modular model provides an absolutely novel *in silico* platform to predict molecular responses of the human skeletal muscle cells to diverse modes of the exercise on three hierarchical levels (metabolic, signaling and gene expression), experimental precise measurements of which are currently methodologically limited or even remain elusive.

In the current state, the model is suitable for testing the plausibility of some physiological hypotheses. For example, the existence of intermediate X factor regulating expression of *PPARGC1A* gene as the example of delayed response gene in human skeletal muscle has been numerically investigated using different versions of the model: considering direct regulation via CREB-like factor or taking into account X factor regulatory role as an intermediate activator of *PPARGC1A* expression.

### Model constraints and further ways for the development

Despite the complexity of the developed modular model, the current version does not consider the influence of many system factors like a hormonal regulation, the influence of processes in the central nervous system, exercise-induced temperature drift in skeletal muscle that hampers precise quantitative reproduction of abrupt changes on different physiological levels on initial stages of the physical exercise. It can be overcomed by means of significant modifications on the muscle fiber recruitment model in order to simulate the transient process due to exercise. Some other constraints described in details below.

GS activity is regulated through multiple mechanisms, including feedbacks mediated by glycogen, blood glucose concentration, rate of glucose uptake, insulin, epinephrine and GS phosphorylation state (Jensen & Lai, 2009; Prats et al., 2009; Jensen et al., 2012b; Palm et al., 2013). However, in the current model GS activity depends on glycogen content only. In our model the post-exercise glycogen synthesis is lower than estimated in the majority of studies (Casey et al., 1995; Jentjens & Jeukendrup, 2003; Burke et al., 2017) because many factors are omitted, such as feeding and associated rises in blood glucose concentration, rate of glucose uptake, sensitivity to and changes in insulin, etc. At the same time, in our model glycogen synthesis is higher than observed during exercise recovery in a fasted state (Maehlum & Hermansen, 1978).

Additionally, our model does not take into account an effect of the muscle fatigue related with the decline in power generated by type II muscle fibres and recruitment of new type II fibres as well as with depletion of the muscle glycogen depot and other substrates. This may play an important role for simulation of moderate and high intensive and/or long-lasting exercise. These limitations provide a direction for the model improvements and should be considered in further works.

Furthermore, the modular nature of the presented model allows the introduction of multiple positive and negative feedbacks between different considered levels: for instance, an impact of kinases altering the activity of enzymes which catalyze reactions of the glycolysis, TCA cycle and fatty acid oxidation in skeletal muscle (Egan & Zierath, 2013, Mounier et al., 2015), Ca^2+^-dependent enhancement of glycolytic enzyme activity and mitochondrial respiration (Gehlert et al., 2015), PGC1α-dependent regulation of the expression of genes encoding glycolysis and malate-aspartate shuttle enzymes (Agudelo et al., 2019). However, all the above-mentioned constraints of the integrated model, rather, present a roadmap for further development and extensions and have methodological solutions via the modular modelling approach used in the study.

### Conclusions

We developed, for the first time, the integrated model of human skeletal muscle incorporating metabolic, signaling and gene expression modules. The model enables to simulate the most important exercise-related signaling (Ca^2+^ and AMPK-related signaling) and RNA expression of early response genes (as a result of activation of transcription factors existing in the cell), as well as expression of delayed response genes (as a result of expression of intermediate transcription factors induced immediately after an exercise). The molecular response of skeletal muscle to contractile activity related to the great number of signaling molecules and genes. The modular nature of the model enables to add new variables and modules thereby increasing both complexity and quality of the model.

## Availability

http://muscle.biouml.org/index.php/Data_availability_for_the_manuscript We have created special wiki page that provides links to:

- GitLab project that contains all source code for the developed model and its modules;
- project on BioUML server which displays the developed model and its modules in visual form as well as provides possibilities for editing and simulation;
- Jupyter notebook (Davies et al., 2020) that provides reproducibility of results presented in the article. The notebook can be started on the BioUML server as well as using Binder technology.

## Acknowledgements

The study has been financially supported by RFBR grants (№ 17-00-00308(K): 17-00-00296, 17-00-00242).

## Author contributions

IA, AV, DP and FK conceived and designed the study. IK and IR contributed to the model reconstruction. IK, IA, AV, SP and FA contributed tools and methods. IA performed the simulations. IK, IA, AV, SP, DP and FK wrote the source code and analyzed the data. AV, SP and RS performed the statistical analysis. IA, VA, OV, DP and FK wrote the manuscript. DP and FK supervised the project. All authors edited the manuscript.

## Supplementary materials

**Supplementary Fig. 1.**
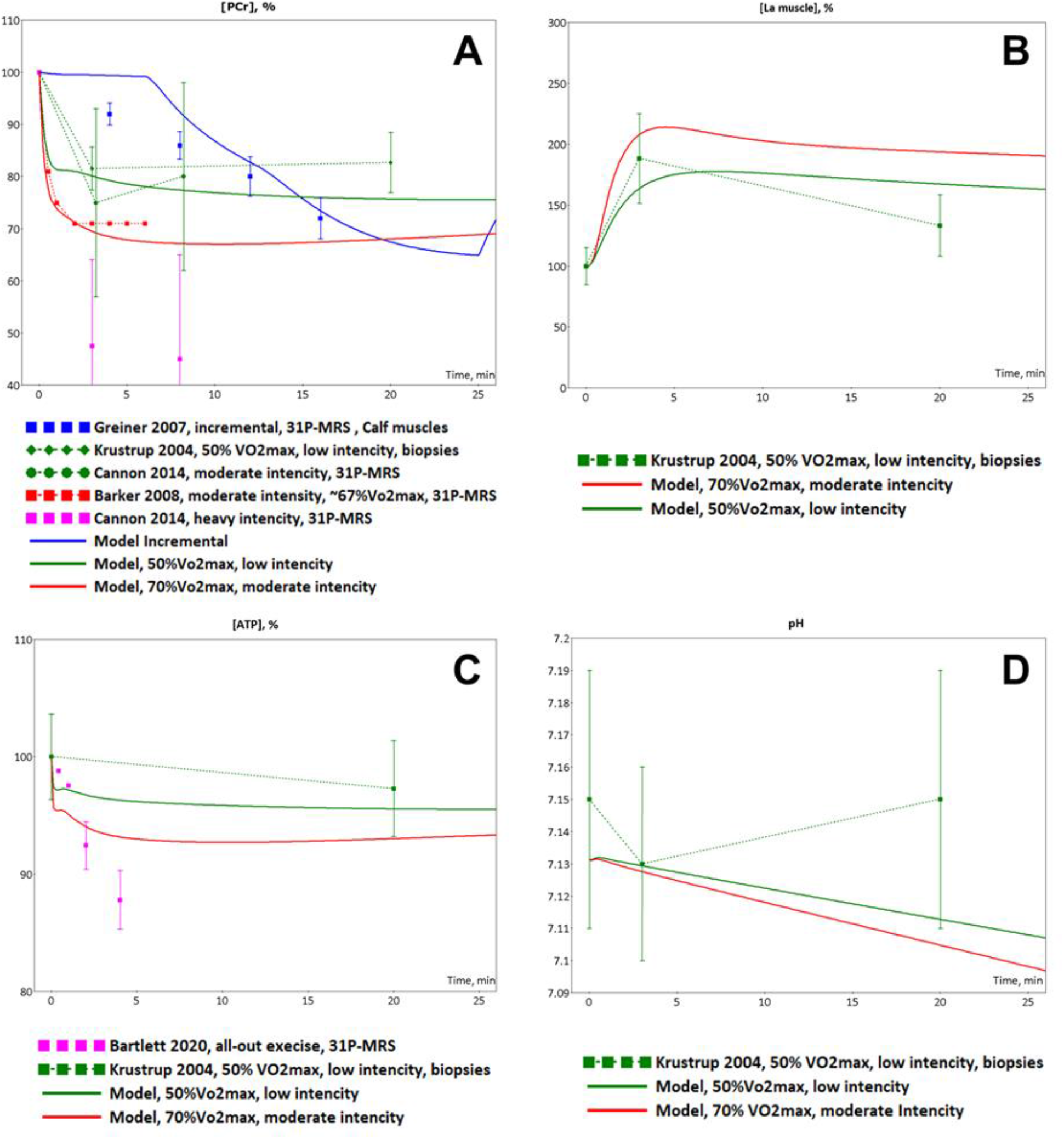
Model validation. The model simulations were compared to experimental data. (A) Model-predicted dynamic changes of PCr concentration in the muscle during incremental (blue line), low (green line) and moderate intensity (red line) exercises (axis X - time of the exercise in minutes) and experimental measurements of PCr concentration changes during incremental (blue square dots, Greiner et al., 2007), low-intensity (green diamond dots (Krustrup et al., 2004), green circle dots (Cannon et al., 2014)), moderate intensity (red square dots (Barker et al., 2008)) and high-intensity (magenta square dots (Cannon et al., 2014)) exercises; (B) Model-predicted dynamic changes of lactate concentration in the muscle during low (green line) and moderate intensity (red line) exercises (axis X - time of the exercise in minutes) and experimental measurements of lactate concentration changes during low-intensity (green square dots (Krustrup et al., 2004)) exercise; (C) Model-predicted dynamic changes of ATP concentration in the muscle during low (green line) and moderate intensity (red line) exercises (axis X - time of the exercise in minutes) and experimental measurements of ATP concentration changes during low-intensity exercise (green square dots (Krustrup et al., 2004)) and all-out exercise (magenta square dots (Bartlett et al., 2020)). The set of experimental data for ATP demonstrates lower (Krustrup et al., 2004) and upper (Bartlett et al., 2020) boundaries for the concentration; (D) Model-predicted dynamic changes of pH in the muscle during low (green line) and moderate intensity (red line) exercises (axis X - time of the exercise in minutes) and experimental measurements of ATP concentration changes during low-intensity (green square dots (Krustrup et al., 2004)) exercise. Experimental data are shown as mean ± SD.

